# Protein Language Models are Accidental Taxonomists

**DOI:** 10.1101/2025.10.07.681002

**Authors:** Logan Hallee, Tamar Peleg, Nikolaos Rafailidis, Jason P. Gleghorn

**Affiliations:** Center for Bioinformatics and Computational Biology, University of Delaware; Synthyra, PBLLC; Department of Medical and Molecular Sciences, University of Delaware; Department of Biomedical Engineering, University of Delaware

**Keywords:** Protein Language Modeling, Protein-Protein Interactions, Taxonomy, Phylogenetics, Negative sampling, Confounders

## Abstract

Protein-protein interactions (PPIs) are fundamental to nearly all biological processes, yet their experimental characterization remains costly and time-consuming. While computational methods, particularly those using protein language models (pLMs), offer higher-throughput solutions, they often report unexpectedly high performance on multi-species datasets. Here, we introduce the accidental taxonomist hypothesis, proposing that neural networks can exploit the phylogenetic distances across labels in protein datasets rather than genuine interaction features. We show that in PPI datasets with random negative sampling, protein pairs for real PPIs are almost exclusively from the same species, while negatives almost always originate from different species. We then demonstrate that pLM embeddings can be used to accurately distinguish whether two proteins share a taxonomic origin, allowing models to “cheat” by learning phylogeny instead of genuine PPI features. By employing a strategic sampling strategy that restricts negative examples to protein pairs from the same species, we reveal a marked drop in model performance, confirming our hypothesis. Compellingly, these strategically trained models still outperform single-species models, suggesting that multi-species data can improve performance if carefully curated. These findings suggest that accidental taxonomist behavior is a particularly influential confounder for PPI, and it is also broadly applicable to any supervised-learning protein dataset.

## Introduction

Proteins are ubiquitous macromolecules that drive biochemical reactions, compose biological structures, and enable physiological signaling, all from organized but straightforward physicochemical forces (1–11). They often accomplish these vital tasks through intricate cross-talk with each other, known as Protein-Protein Interactions (PPIs). In its most basic form, a PPI involves two proteins exhibiting physical contact that mediates chemical or conformational change, especially with non-generic function (12–14). At scale, paired PPIs can form complex networks and pathways, and are central to life sciences due to their implications in studying fundamental biology, disease, and potential therapeutics. However, traditional characterization of PPIs through *in vitro* and *in vivo* methods is expensive and time-consuming. Established higher-throughput methods like Yeast 2-Hybrid (Y2H) can identify thousands of PPIs once libraries are prepared at relatively inexpensive costs. More recent methods, such as Cell-Free Two-Hybrid (CF2H), enable the coupling of cell-free protein expression and precise detection capabilities that can correlate to binding affinity (15, 16). Despite these advances, the scale of the interactome (over 200 million pairwise PPIs in humans) necessitates a major improvement in throughput. A comprehensive, computable, dynamic, and accurate interactome could serve as the base blueprint of cellular function. To this end, many groups are researching computational methods to predict PPIs from protein identifiers (17, 18). Some methods use protein or gene IDs to search and extrapolate from established databases, others leverage protein structure solutions or predictions to infer PPIs, and many recent methods leverage Protein Language Models (pLMs) to infer PPIs from primary sequences (14, 19–22). These platforms, particularly pLMs, offer an attractive solution with the possibility for massively high-throughput screening, amino acid- or atom-level analysis of PPI mechanisms, and inference of functionality (18).

pLMs have shown various levels of claimed success in predicting PPI. However, as with any machine learning model, there are many possible biases in dataset composition that can lead to inflated model performance. PPIs are an interesting machine learning and data compilation problem; while numerous PPI databases curated from the literature contain hundreds of thousands of real PPIs (and millions of likely ones), PPIs are extremely dependent on the biochemical context of the experiment. Factors such as post-transcriptional and post-translational modifications (PTMs), chaperone-mediated folding, protein cofactors, mutagenesis, pH, and many others ensure that recorded PPIs between two entities are rarely *always* true positives. Because of this, we reason that PPI machine learning models trained on general PPI data actually learn the relative plausibility of PPI, without accounting for biochemical context. In other words, we reason that pLM-based methods may infer that there exists a combination of biochemical factors, like PTMs and allosteric changes, that allow for a PPI without guaranteeing that the combination of characteristics ever occurs in reality. Therefore, positive PPI data and prediction come with a high level of ambiguity. Additionally, it is impossible to prove empirically that something cannot happen, so there are no “real” negative PPI examples to contrast against PPI positives. The Negatome project has improved the availability of extremely unlikely PPIs through manual analysis; however, even then, all of Negatome-2.0 data splits include only 6,000 negative PPI protein pairs when combined (23). This extreme data imbalance ensures that any binary prediction machine learning model trained on the unmodified available data will almost always predict “positive” (14, 24). Previous popular negative curation methods leveraged information about protein subcellular compartment localization to estimate negatives, under the logic that proteins that localize to different cellular regions are unlikely to interact (25). While this logic is reasonable, proteins that do not interact *in vivo* due to their location may be able to interact *in vitro*, which is still important information in the coming age of protein design (14, 26). In our earlier work around SYNTER-ACT, we posited that pLMs are *accidental localizers*: pLM features can be used to predict the subcellular compartment localization of proteins from their primary sequence with high accuracy (27–29). Thus, pLM-based PPI models trained on real positives and subcellular negatives “cheat” and learn to predict the difference in localization, rather than PPI (or in addition to PPI) (14, 30).

More recently, papers by Bernett et al. established a “goldstandard” dataset built with different sequence-similarity-based biases in mind, attempting to control for pLMs memorizing domains or motifs (20). Their balanced binary PPI dataset was constructed using positives from the Human Integrated Protein-Protein Interaction Reference (HIPPIE), and negatives generated through randomly selecting protein pairs to match the number of positives (20). This random selection method for PPI negatives is currently the most popular approach, which leverages the assumption that randomly generated or paired amino acid sequences should not induce any meaningful PPI when expressed together; a good assumption when considering the highly specific nature of many PPIs. It also, at least partially, alleviates the accidental localizer issue by ensuring a mix of subcellular localization trends in the negative examples. Therefore, a model cannot purely rely on the localization confounder. Evaluation splits of the Bernett dataset were strategically partitioned so that there were no node degree biases, no sequence overlap (referred to as C3 by Park and Marcotte (31)), and a maximum sequence similarity of 40% between any of the train, validation, and test sequences. Works from Ko et al. and Bernett et al. have benchmarked many established pLM- or neural network-based PPI methods on this gold-standard dataset and have shown that performance is roughly limited to 0.7 AUC and 65% accuracy with current modeling capabilities (21, 22, 32, 33).

Yet, we make a simple observation that on multi-species datasets, unlike Bernett’s gold-standard *human-protein-only* PPI dataset, pLM-based methods showcase <u>much</u> higher performance metrics on withheld data. Even our SYNTERACT model, trained on a modified version of BioGRID, showcased over 90% accuracy on our withheld test set (14, 34). In Ko et al.’s paper, we also observe high AUC scores ranging from 0.8 to over 0.9 with multi-species performance (STRING-db v11, C3, 40% CD-HIT) (22). We believe this trend holds for much of the literature (25, 33, 35–40). Of course, to a certain degree, this is expected, as more species means more data and sequence diversity, and machine learning models should improve accordingly. However, the performance difference is staggering.

In this paper, we highlight observations and experiments that explore PPI performance discrepancies and put forward a hypothesis for the suspiciously high multi-species PPI performance reported in the literature. Specifically, we expand upon work analyzing the encoded phylogenetic relationships in pLM embeddings by demonstrating that neural networks trained on pLM embeddings can predict the taxonomic origin of protein primary sequences, even at extreme levels of discrimination (41–43). We further postulate and support that it should be even easier than predicting the precise taxonomic classification of one protein to distinguish whether two proteins share a taxonomic origin. If it is possible for pLM-based methods to distinguish a shared taxonomic origin from two protein sequences, it implies that PPI models trained on multi-species data could be inadvertently learning taxonomic features and not PPI. We refer to this hypothesis as the “**accidental taxonomist**,” where multi-species PPI models exploit the phylogenetic distance of negative examples to artificially inflate their performance. We further support our hypothesis by training a variety of pLM-based models in two conditions, where (1) the negatives are sampled randomly from BioGRID and (2) the negatives are sampled randomly from *within the same species* of BioGRID data. We show on rigorously split evaluation sets, similar to Bernett’s goldstandard (C3, 40% CD-HIT), that option (1) will “cheat” and option (2) cannot; implying a straightforward solution to this problem through strategic sampling.

Unfortunately, while we uncover the accidental taxonomist for PPI prediction, it is also potentially a key confounder in any supervised protein dataset, where certain classes or sections of labels may belong to a phylogenetic region, thereby compromising the underlying performance of the model despite reasonable evaluation set metrics. We intend for researchers to carefully consider phylogenetic distances and taxonomic biases to establish standardized controls that can be implemented to improve the performance and utility of pLM-based supervised learning.

## Results

### Multi-species PPI datasets are subject to inherent phylogenetic biases

After shuffling a common multi-species PPI data source, BioGRID (34), we observed that the randomly paired examples only shared the same organism of origin about 31% of the time on average (**Figure 1**). Assuming a deep learning model could predict taxonomic origin perfectly, it would predict the positive examples of a PPI dataset compiled this way nearly 100% correctly, as the vast majority of real PPIs come from sequences of the same species, and would predict negative examples with approximately 69% accuracy. The performance metrics of this scenario on a balanced dataset would yield 85% accuracy, an F1 score of 0.87, an MCC of 0.73, and an AUROC of 0.85.

**Fig. 1.**
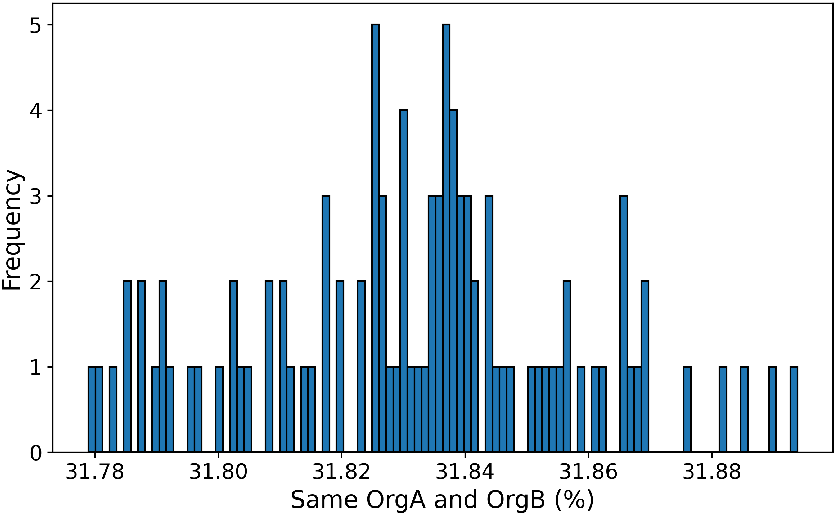
Histogram of the percentage of rows in BioGRID that maintain matching species after shuffling. Shuffling was repeated 100 times with different seeds to demonstrate the consistency of this finding.

### pLM features can be used to predict precise taxonomic origin

We benchmarked several popular pLMs on large protein datasets with labels based on taxonomic discrimination, ranging from domain to species. Datasets were embedded and fed to linear probes, highlighting the intrinsic correlation between the embeddings and the tasks. We also trained on a similar balanced binary classification dataset where pairs of sequences were either from the same species or not. Performance on these compiled datasets is showcased in **Figure 2**.

**Fig. 2.**
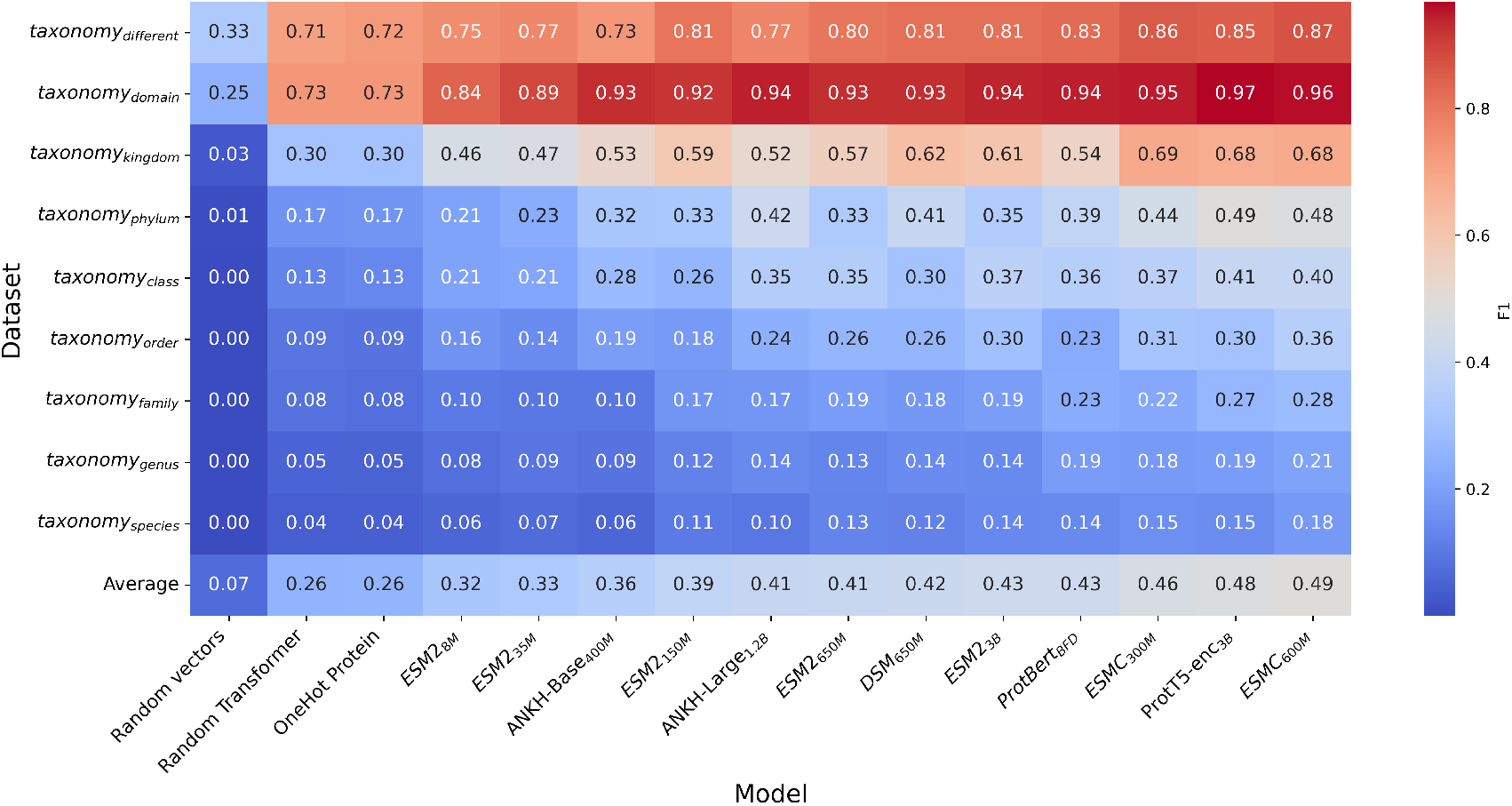
Performance heatmap of all models and datasets evaluated comparing pLM embedding correlations to taxonomic discrimination. Weighted F1 scores are presented for each model-dataset combination, chosen because of the large class imbalance. The different datasets constructed by taxonomy are arranged along the rows, with individual model performance within each column.

**Fig. 3.**
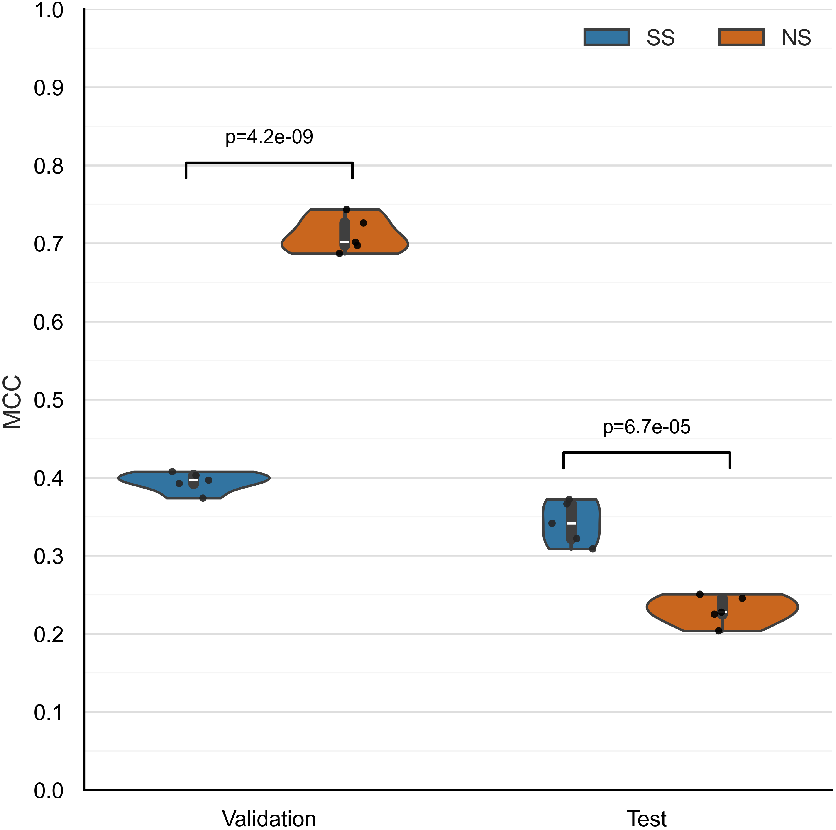
Comparison of SS and NS approaches with Violin plots of MCC scores on the validation and test sets over five PPI training runs. The five NS and SS score distribution differences are highly statistically significant.

**Fig. 4.**
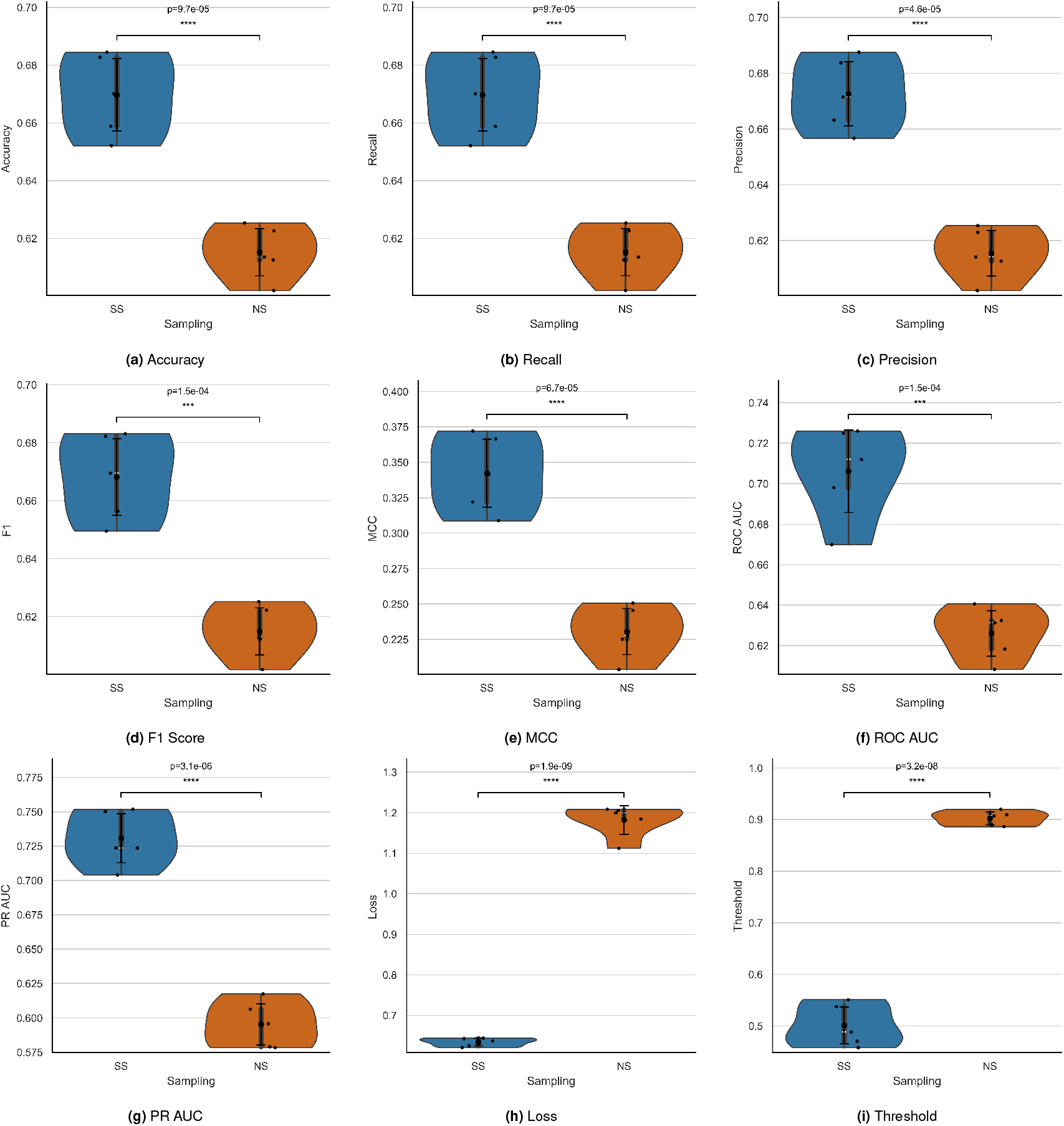
Comparison of SS and NS approaches across various measured performance metrics on the test set. SS is shown on the left of each plot in blue, with NS on the right in orange.

The random vector column of **Figure 2** showcases truly random performance on each test dataset, which varies significantly based on the number of classes in each dataset. A transformer with random weight initialization and one-hot encoding demonstrated basic performance gains from the addition of basic sequence features alone, also known as a homology controls (29). Every other probe was fed embeddings from a pLM that had undergone significant semi-supervised denoising, which clearly enhanced the feature vectors for taxonomic classification, as evidenced by the test F1 scores compared to random and homology controls.

Highlighted in **Figure 2**, most pLM embeddings could be used to almost perfectly predict sequences with vast phylogenetic distances at the domain level in taxonomy_*domain*_ (corresponding to Archaea, Bacteria, and Eukarya), with the best performer at 0.97 F1 (ProtT5). At the kingdom level, taxonomy_*kingdom*_, this performance dropped to 0.69 F1 (ESMC_300*M*_), 0.49 F1 (ProtT5) at phylum, 0.36 at order (ESMC_600*M*_), 0.28 at family (ESMC_600*M*_), and 0.21 at genus (ESMC_600*M*_). At the most discriminative level, taxonomy_*species*_ (among 618 species classes), the best-performing pLM embeddings yielded a significantly betterthan-random performance, with an F1 score of 0.18, compared to 0.00 F1.

The model probes performed much better on taxonomy_*different*_, identifying if two input sequences share a species or not, compared to precise species classification in taxonomy_*species*_. Most pLMs exceeded 0.8 F1 on this task, with ESMC_600*M*_ embeddings yielding the best F1 score of 0.87.

### Strategic sampling prevents “cheating” on PPI sets

We attempted to rectify the accidental taxonomist behavior by never feeding a “negative” PPI to a model from different species. Instead, we randomly selected sequence pairs from within the same species to construct negatives; we refer to this approach as *Strategic Sampling (SS)*. This simple change from the *Normal Sampling (NS)* approach, where negatives are sampled randomly from the entire multi-species dataset, resulted in vastly different performance metrics on the validation and test sets. The validation sets follow the same sampling as the training set, but both test sets follow the SS approach to trick the NS model if it has started to cheat. The NS models averaged a 0.71 MCC on the validation set, compared to a 0.39 MCC for the SS version, implying that the NS models heavily exploit phylogenetic distance measurements. This is also evident in the test sets, where the NS model achieved a 0.23 MCC, compared to 0.34 MCC for the SS models; a significantly lower performance from the NS model on the SS test set, as it is unable to effectively exploit phylogenetic distances. Both differences are statistically significant, with a *t*-test *p <* 0.0001 for each (*n* = 5). Notably, the 0.23 MCC score from NS is much better than random performance (equivalent to 62% accuracy, 0.61 F1 score, and 0.63 AUROC), meaning that the NS models do not solely rely on phylogenetic distances and appear to still learn aspects of PPI. Excitingly, the best test SS MCC of 0.37 is approximately 23% higher than the current best MCC scores on Bernett’s dataset (0.30 MCC), despite similar dataset construction rigor. This implies that incorporating multi-species data is an effective way to enhance performance and generalizability when controlled for correctly.

## Discussion

Computational PPI prediction is a vital problem to advance due to its widespread applicability across life science domains. With performance plateauing around 65% accuracy and 0.3 MCC on rigorously constructed single-species PPI datasets (20, 21), and multi-species PPI datasets showcasing near-perfect performance, we aimed to explore the underlying cause of this inflated performance. We noticed that multi-species datasets such as BioGRID, and random negative sampling exemplify this data composition bias (19, 34, 44). If you shuffle the paired sequence A and sequence B columns in a dataset derived from BioGRID, only 30% of those generated negatives share a species of origin. This means that a model that can *perfectly* distinguish the taxonomic origin of a protein sequence would achieve excellent classification metrics of roughly 85% accuracy, 0.87 F1, an MCC of 0.73, or AUROC of 0.85 on a balanced binary PPI dataset, *without learning anything about PPI*. Of course, perfectly distinguishing taxonomic origin and learning PPI features would have even higher performance. These observations closely align with reported multi-species results from the literature (22, 25, 33, 36–40). It is worth noting that BioGRID contains over 44% human data, with other large percentages coming from model organisms. Much larger and more diverse repositories, like STRING-db, would exacerbate this issue further, with an even smaller fraction of protein pairs coming from the same species if the entire dataset were shuffled.

To explore the capabilities of pLM-based methods to distinguish the taxonomic origin of primary sequences, we expanded upon existing work (41, 42) on pLM embeddings and encoded phylogenetic distances to characterize how well various neural network probes can classify taxonomic origin across a wide range of discrimination, from broad domain to species. Unsurprisingly, the best-performing model on average, ESMC_600*M*_ (45, 46), achieved almost perfect performance in predicting sequences with vast phylogenetic distances at the domain level in taxonomy_*domain*_ (corresponding to Archaea, Bacteria, and Eukarya). We observed a gradual decline in overall performance as the phylogenetic distance of the labels decreased. Interestingly, at the most discriminative label set in taxonomy_*species*_, among 618 species classes, ESMC_600*M*_ maintained a much better-than-random performance of 0.18 F1 score (vs. 0.00 random), and sometimes the exact species of origin could be determined from only a single primary sequence. Considering the training dataset utilized sequences that were maximally 40% similar to the validation and test sets, we were surprised to achieve better-than-random performance at the species level and encourage future work to explore which specific combination of learned features from semi-supervised denoising drives this performance. Nevertheless, despite being better than random, a 0.18 F1 score is far from reliable performance.

However, in line with our hypothesis, we demonstrated that when fed two sequences in taxonomy_*different*_, aligned with the standard workflow of PPI prediction, neural networks could effectively predict if each sequence came from a different species. In fact, most pLMs tested exceeded 0.8 F1 score on this balanced binary prediction task, with ESMC_600*M*_ achieving 0.87 F1. Even the homology controls exceed 0.7 F1. Thus, even though neural networks cannot predict the exact species from pLM embeddings, they can reliably detect whether two sequences have a nonzero phylogenetic distance. We refer to this finding as the accidental taxonomist, where pLM-based methods exploit phylogenetic distance distributions across classes to inflate their performance. We reasoned this is particularly applicable to PPI datasets, where typical datasets have pairs labeled “1” with no phylogenetic distance and most pairs labeled “0” with some distance.

We attempted to remedy the inflation in multi-species PPI performance by comparing normal sampling (NS) to strategic sampling (SS). Instead of building negatives from randomly assigned protein pairs, we restricted the assignment to sequences within the same species. The results are clear: across five different NS and SS large-scale PPI runs, covering over 4.5 million high-quality datapoints with rigorous evaluation splits, NS models perform exceptionally well on NS data splits while failing on SS splits. This large discrepancy of 0.7 MCC vs. 0.2 MCC highlights the difference in performance when models exploit phylogenetic distances versus when they cannot. In contrast, SS models maintained consistent performance on SS validation and test sets, achieving an MCC of approximately 0.4 on both. Importantly, despite comparable dataset construction rigor, our multi-species SS PPI models all exceed the 0.3 MCC plateau for single-species PPI, implying that adding sequence diversity continues to improve PPI performance and generalizability when properly controlled. More reported metrics are shown in **Supplemental Figure 4**.

From this work, our previous identification of accidental localizers, and the outstanding contributions of Bernett, Ko, Rost, and many others, exploring the flaws and biases behind PPI prediction, we can safely assume that very high PPI performance metrics (for example 0.9 AUC, 0.5 MCC, F1, 80% accuracy, etc.) are likely due to some form of data leakage or underlying confounders, in addition to genuine learning of PPI. High-throughput computational PPI remains an unsolved problem, and dataset construction and curation challenges remain major obstacles.

Unfortunately, our analysis underscores a more fundamental flaw in pLM dataset compilation beyond PPI prediction. For example, one common application of pLM embeddings is predicting Enzyme Commission (EC) numbers, a categorization scheme for different enzymatic reactions. If a given EC number occurs only in a specific taxonomic family, a model trained on such data may inadvertently learn to classify by taxonomy rather than by enzymatic function. This same principle applies to any supervised protein-learning task, including regression, where a specific label range could belong exclusively to a taxonomic rank.

It is conceptually intuitive to imagine an attention-based neural network exploiting confounders during multiple-input tasks, such as PPI, as these models directly compare two inputs via dot product (47). However, even simpler neural networks and single-protein input datasets are not immune to the accidental taxonomist effect. Neural networks are capable of storing various training distributions into their weights via gradient descent (48–50). For instance, a simple feed-forward network could organize one layer of weights to correspond to the average taxonomic features for each class, causing inputs with specific matching taxonomic features to produce higher values after this layer.

Additionally, we suspect that models leveraging DNA or codon vocabularies to study proteins will be even more susceptible to the accidental taxonomist phenomenon. Codon usage bias is a powerful feature for predicting taxonomic origin on its own (43, 51), adding another layer of abstraction on top of amino acid sequences for neural networks to exploit during training.

Finally, there may be other “accidental” phenomena where unintended physical, chemical, biological, or social trends, surrounding which proteins researchers choose to study, distract machine learning models from learning the biological principles of interest. We hope to see future work exploring how neural networks “cheat” on protein datasets and how improved dataset design and controls can mitigate these confounders.

## Methods

### Prototyping phylogenetic bias impact

The February 2025 release of BioGRID (4.4.243) was downloaded, and protein identifiers were mapped to UniProt accessions, with Swiss-Prot prioritized. IDs were mapped to sequences using the UniProt ID mapper tool https://www.uniprot.org/id-mapping and the primary se-quence and organism of origin were recorded for each protein pair (34, 52). The six columns in our dataset include each UniProt ID (A, B), each sequence (SeqA, SeqB), and each organism (OrgA, OrgB). To simulate negative pair generation, we shuffled the OrgA and OrgB columns 100 times and summed the number of occurrences of the same organism, as shown in **Figure 1**. We simulated the classification metrics for a perfect OrgA = OrgB vs. OrgA ≠ OrgB classifier by building random predictions and labels. The 100,000 data points consisted of 50% labeled one and 50% labeled zero, where 31% of the zeros were false negatives per the results of the BioGRID shuffling. The actual predicted probabilities were randomly generated from a uniform distribution between 0.51 and 1.00 for true positives, 0.00 to 0.49 for true negatives, and 0.51 to 1.00 for false negatives, allowing for the estimation of the AUROC. Accuracy, precision, recall, F1, and MCC were also calculated.

### Probing taxonomic features

Datasets at different taxonomic discriminations were constructed from Swiss-Prot, downloaded on 7/22/2025 (52). We kept the taxonomic lineage column to build labels and retained sequences of lengths greater than 20 and less than 2048. Sequences were deduplicated and clustered at 40% sequence identity with a word size of two using CD-HIT (53), and clusters were randomly assigned to validation and test splits until their count exceeded 10,000 entries, respectively. For each taxonomic discrimination we only kept labels that had at least 100 examples total, resulting in three domain, 17 kingdom, 68 phylum, 126 class, 248 order, 371 family, 545 genus, and 618 species classes. The total training set sizes consisted of 444k domain, 450k kingdom, 458k phylum, 451k class, 446k order, 434k family, 419k genus, and 265k species examples. We enforced that there was no sequence or cluster overlap between any splits to ensure evaluation rigor. The datasets are referred to as taxonomy_*domain*_, taxonomy_*kingdom*_, taxonomy_*phylum*_, taxonomy_*class*_, taxonomy_*order*_, taxonomy_*family*_, taxonomy_*genus*_, and taxonomy_*species*_ throughout the work for clarity of the discrimination level.

The taxonomy_*different*_ dataset was compiled by taking taxonomy_*species*_ and randomly constructing protein pairs from within each data split. 50,000 random sequences from the same species were sampled for the training split, 500 for validation, and 500 for testing. Then, the dataset was balanced by randomly selecting sequences from different species. The final dataset, comprising 100,000 training, 1,000 validation, and 1,000 test sequences, presented a binary prediction problem: determining if two sequences had any phylogenetic distance between them, where protein pairs were labeled as zero if they were from the same species and one otherwise.

Models and datasets were probed using the **Protify** package, an open source project aimed at standardized, effective, and high-throughput analyses with chemical language models (54). We embedded all datasets using mean and variance pooling on the last hidden state of various pLMs, resulting in a consistently sized vector for each protein describing the distribution of the models’ last hidden state. For taxonomy_*different*_, which took two sequence inputs per example, those pooled vectors were stacked together. We chose many popular pLMs for analysis, including ESM2 eight million parameter version (ESM2_8*M*_), ESM2_35*M*_, ESM2_150*M*_, ESM2_650*M*_, ESM2 three billion parameter version (ESM2_3*B*_), ESMC_300*M*_, ESMC_600*M*_, DSM_650*M*_, ProtBERT (big fantastic dataset version), ProtT5-enc_3*B*_ (ProtT5), ANKH-Base_400*M*_, and ANKH-Large_1.2*B*_ (45, 54–57). For the T5-based models, ANKH and ProtT5, only the encoders were used for embedding generation. Three controls were also used: random vectors, which embed each protein with a randomly generated vector (*x* ∼𝒩 (0, 1)); random transformer, which embeds proteins through a re-initialized equivalent of ESM2_35*M*_ ; and OneHot protein, which replaces the pLM embeddings with one-hot encoded strings (54).

Within Protify, linear neural network probes were trained with the architecture:

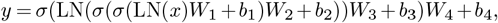

mapping input *x* ∈ ℝ^*d*^ to output *y* ∈ ℝ^*o*^ where *o* was the number of classes in the dataset. *W*_1_ ∈ ℝ^*d×h*^, *W*_2_ ∈ ℝ^*h×h*^, *W*_3_ ∈ ℝ^*h×p*^, *W*_4_ ∈ ℝ^*p×o*^, *b* corresponded to the associated bias term in the linear layer, *σ* was ReLU (58), *h* was 8,192, *p* was calculated with

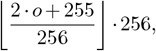

and a dropout of 20% was applied before every MatMul, except the first one (59). We optimized cross-entropy loss with a batch size of 64, learning rate of 1*e*^−4^, AdamW optimizer, weight decay of 0.01, and evaluation on the validation set until a patience of five was exceeded, comparing the validation loss. The best weights were kept and evaluated on the test set.

### Strategic sampling experiment

The protein sequences in BioGRID 4.4.243 were deduplicated and clustered at 40% sequence identity with a word size of two using CD-HIT (53). Clusters were added to a list until their paired sequences exceeded 5,000 BioGRID examples for validation and testing, respectively. We ensured that every data split was disjoint in terms of sequence and cluster ID, following the C3 strategy from Park and Marcotte (31). Negatives were generated within each data split by either sampling random proteins from the clusters (Normal Sampling / NS) or sampling random proteins from the clusters (Strategic Sampling / SS) that are from the same organism. If a negative pair was found in the positive pairs we discarded it and sampled again. The final datasets for NS and SS both consisted of 4,523,432 training examples, 10,070 validation examples, and 10,034 test examples, with a perfectly balanced label distribution; the organism distributions between the positive and negative examples were approximately the same, and the datasets used a total of 76,045 unique protein sequences.

We trained a custom hierarchical transformer neural network to map pLM embeddings to binary logits. The embedding model of choice was ESMC_600*M*_ (45, 46), which we froze and extracted the last hidden state for each sequence, truncating it to a length of 512 to minimize computational expense. This followed recent findings that larger maximum lengths have a minimal effect on PPI performance (60). Because these are not intended as production models and are used for comparison between NS and SS, this truncation was acceptable.

For each sequence A and B, there was an identical track of a linear layer projections from ESMC_600*M*_ hidden size of 1,152 to 512, one transformer block with hidden size 512, and then token-parameter cross-attention which pooled every protein sequence into consistently sized matrices 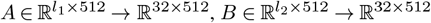. *A* and *B* were then concatenated into *X* ∈ ℝ^64*×*512^, which served as the input to the rest of the model. The remainder of the model consisted of four transformer blocks, interleaved with linear layers, which reduced the dimensionality from ℝ^64*×*512^ to ℝ^64*×*64^, projecting down by a factor of two at every transformer block. The final ℝ^64*×*64^ matrix was mean-pooled into a ℝ^64^ vector and fed to a final linear layer to build the logits ℝ^1^. An expansion ratio of 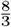 was used for all transformer block MLPs and all linear layers (aside from attention layers) used a dropout of 0.1. Binary cross-entropy was employed alongside the AdamW optimizer (63), a batch size of 128, learning rate of 1*e*^−4^, evaluation on the validation set every 5,000 steps, and all models were trained for one epoch on a single GH200 GPU; the weights with the best validation MCC were kept and test scores were reported.

In case each model curated slightly different decision boundaries, and different thresholds were optimal to split logits into positives and negatives, we split the precision-recall curve into 100 discrete points and tried each threshold. The threshold used for the metrics calculations was chosen based on which one produced the maximal F1 score. Technically, macro F1 was used for threshold selection and weighted averages were reported for F1, precision, and recall (**Supplemental Figure 4**); however, because the datasets were balanced, this did not impact the final result.

## DATA AND RESOURCE AVAILABILITY

Selected datasets, code, and model weights will be available at github.com/Gleghorn-Lab/PLMConfounders. Protify can be accessed here: github.com/Gleghorn-Lab/Protify.

## AUTHOR CONTRIBUTIONS

Conceptualization (LH, JPG), Model architecture (LH), Data Curation (LH, TP), Investigation (LH, TP), Formal Analysis (LH, TP, NR), Writing – Original Draft (LH), Writing – Review & Editing (LH, TP, NR, JPG), Supervision (LH, JPG), Project Administration (JPG), Funding acquisition (LH, JPG).

## ACKNOWLEDGEMENTS

The authors thank Katherine M. Nelson, Ph.D., for reviewing and commenting on drafts of the manuscript. This work was partly supported by the University of Delaware Graduate College through the Unidel Distinguished Graduate Scholar Award (LH), the National Science Foundation through NAIRR pilot 240064 (JPG), and the National Institutes of Health through NIGMS T32GM142603 (LH, NR), R01HL178817 (JPG), R01HL133163 (JPG), and R01HL145147 (JPG).

## Supplementary Material

